# Understanding the Genetics of Viral Drug Resistance by Integrating Clinical Data and Mining of the Scientific Literature

**DOI:** 10.1101/2021.10.26.465417

**Authors:** An Goto, Raul Rodriguez-Esteban, Sebastian H. Scharf, Garrett M. Morris

## Abstract

Drug resistance caused by mutations is a public health threat for existing and emerging viral diseases. A wealth of evidence about these mutations and their clinically-associated phenotypes is scattered across the literature, but a comprehensive perspective is usually lacking. This work aimed to produce a clinically-relevant view for the case of Hepatitis B virus (HBV) mutations by combining a chronic HBV clinical study with a compendium of genetic mutations systematically gathered from the scientific literature. We enriched clinical mutation data by systematically mining 2,472,725 scientific articles from PubMed Central in order to gather information about the HBV mutational landscape. By performing this analysis, we were able to identify mutational hotspots for each HBV genotype (A-E) and gene (C, X, P, S), as well as the location of disulfide bonds associated with these mutations. Through a modelling study, we also identified a mutational position common in both the clinical data and the literature that is located at the binding pocket for a known anti-HBV drug, namely entecavir. The results of this novel approach shows the potential of integrated analyses to assist in the development of new drugs for viral diseases that are more robust to resistance. Such analyses should be of particular interest due to the increasing importance of viral resistance in established and emerging viruses, such as for newly-developed drugs against SARS-CoV-2.

## Introduction

Gene mutations that confer drug resistance to pathogens are an emerging public health and medical threat with, additionally, potential consequences for current and future pandemic response. A step towards combating this scourge is to study the way in which particular mutations lead to viral drug-resistant phenotypes. Viral species hosted by an individual patient can contain multiple mutations that interact to produce specific phenotypes. To understand mutational interactions, the scientific literature can help in interpreting existing knowledge on the biological mechanisms associated with viral mutation-phenotype relationships. Here we explore the feasibility of a novel integrated approach that combines clinical and text mining data to gather existing knowledge and produce new insights on drug resistance based on data from viral species hosted by individual patients, and in particular for the case of HBV. Such approaches have the potential to be applied to other viral diseases, singularly due to the growing acknowledgement of the critical importance of viral mutation monitoring in patients.

Hepatitis B is an infectious disease that affects approximately 292 million people worldwide. Despite being a major global health concern, it has been estimated that only 10% of individuals who are chronically infected with HBV have been diagnosed, and only 5% of those individuals who are eligible for treatment receive an antiviral therapy.^1^ Chronic HBV infection is known to progress to cirrhosis of the liver in up to 40% of untreated patients;^2^ and in 2015, HBV led to an estimated 887,000 deaths, which were largely caused by cirrhosis of the liver and hepatocellular carcinoma (HCC).^3^ There are two classes of treatments that are known to be effective against suppressing HBV infections: interferons and nucleotide analogues. These treatments can reduce the patient’s viral load and the main impacts of the disease on the patient. However, although these treatments have been available for nearly two decades, they have not eliminated HBV.^4^

HBV is classified into ten HBV genotypes, A to J, each with a distinct geographic distribution.^5,6^ The infectious HBV virion, also known as the *Dane particle*, is a spherical, double-shelled structure with a diameter of 42 nm.^7^ This particle consists of an outer lipoprotein envelope and an inner nucleocapsid core, which encapsulates the viral genome.^6^ The genome of HBV is approximately 3.2 kilobase pairs long and has a partially double-stranded circular DNA.^7^ The viral genome codes for all five viral proteins required for HBV replication: the HBV surface antigen; the HBV core antigen; the HBV envelope antigen; the X protein; and the HBV reverse transcriptase/polymerase.

There are four known genes in the viral genome: C, X, P, and S. Gene C encodes the core protein and it also produces the pre-core protein by promoting an upstream AUG start codon as its start codon.^8^ Genes P and S encode for DNA polymerase and HBsAg, respectively. The S gene is an open long reading frame that is divided into three sections based on the start codons: pre-S1, pre-S2, and S.^9^ The mechanism and function of the protein coded by gene X, HBV X antigen, remains largely unknown, but it is known to be associated with the development of liver cancer.^10^

Due to its high mutation rate, with commonly-accepted rates of ~2.0×10^−5^ nucleotide substitutions per site per year,^11^ HBV is able to become resistant to HBV drugs relatively quickly. With the emergence of drug resistance, patients may experience a notable compromise in the efficacy of anti-viral therapy and deterioration of the disease condition.^12,13^ Thus, investigation of resistance mutations is crucial for the successful development of robust HBV drugs. Lamivudine (phosphorylated lamivudine) and adefovir dipivoxil were the first nucleotide analogues that were developed, but had limited success because of the development of resistant variants. Resistance mutations pertaining to nucleotide analogues are usually reported in the polymerase gene of HBV DNA (*i.e*. gene P).^14^ The resistance phenotype is typically a consequence of nucleotide analogues failing to bind to DNA polymerase.^15^

Lamivudine acts as a nucleoside reverse transcriptase inhibitor. In HBV, the active metabolite of lamivudine becomes incorporated into the viral DNA by HBV polymerase which results in DNA chain termination.^16^ The active metabolite of adefovir, adefovir diphosphate, that becomes phosphorylated inhibits HBV reverse transcriptase by competing with the natural substrate deoxyadenosine triphosphate, and similarly causes DNA chain termination after it becomes incorporated into the viral DNA.^17^

After conducting randomized clinical trials of lamivudine, it was found that, compared with placebo, HBV mutant variants were associated with reduced sensitivity to lamivudine in approximately 30% of patients after only a year of treatment.^18,19^ In addition, mutations causing resistance to adefovir dipivoxil were detected in up to 20 to 29% of patients after five years of treatment.^20,21^ However, more-recently developed nucleotide analogues, such as entecavir and tenofovir disoproxil, have been reported to dramatically decrease the rate of development of resistance. Entecavir inhibits all three activities of the HBV polymerase by competing with the natural substrate deoxyguanosine triphosphate by base priming, reverse transcription of the negative strand from the pregenomic messenger RNA, and synthesis of the positive strand of HBV DNA.^22^ In addition, the active metabolite of tenofovir, tenofovir diphosphate, inhibits hepatitis B polymerase by direct binding competition with the natural deoxyribonucleotide substrate. After tenofovir diphosphate becomes integrated into the DNA, it results in viral DNA chain termination.^23^

In the case of entecavir, only 1.2% of patients developed a resistant strain after five years of treatment if they had never been treated with nucleotide analogues previously.^24^ For tenofovir disoproxil, there were no clinically significant resistant variants identified during up to seven years of follow-up.^25^ Cross-resistance is also known to occur and is another concern for the development of drug resistance. For example, between lamivudine- and entecavir-resistant HBV strains, the cumulative probability of developing entecavir resistant variants is more than 50% for those individuals with lack of resistance toward lamivudine.^24^

In order to create new therapies that can circumvent or negate mechanisms of drug resistance, we need to understand resistance mutations better with the aid of analytics tools such as text mining. Text mining has been applied to extract evidence for generic drug resistance, impact of mutations in disease and to harvest viral mutation data.^26–29^ It has also been leveraged to create comprehensive databases of the viral mutation literature.^30,31^ There is no report, however, that it has been used to produce new insights that support clinical data analysis of viral mutations with integration of literature data. This study aimed to build on a clinical study that ultra-deep sequenced HBV quasispecies from a European cohort^32^ by comparing the data from this clinical study against genetic mutation information automatically extracted from the scientific literature regarding all known launched anti-HBV drugs. Such data are typically scattered across many publications; therefore, we built a pipeline to mine literature sources systematically. In order to extract mutations from the literature we applied a rule-based text mining approach.^33–35^ With our approach, we were able to produce a landscape of disease-specific viral mutations associated to drug resistance, which allowed us to derive new insights into mechanisms associated to drug resistance.

There exist manually curated databases^36–39^ that aim to gather HBV resistance mutations from patients’ data. However, it is hard to evaluate their coverage since they do not give a complete list of resistance mutations and instead provide a mean to analyze the genetic sequences that the user has obtained. In addition, most of these databases are not open source. Hence, the use of text mining approaches that are able to mine through the literature efficiently are essential to tackle the challenge of identifying known resistance mutations.

## Methods

### Code and Supplementary Information

The analysis was performed using standard Python packages (*e.g. ElementTree^40^* for parsing XML-formatted literature, *re* for identifying information about mutations, and *pandas* for data analysis). The resulting code and supplementary information (Figures S1-S18) is available at: https://github.com/angoto/HBV_Code.

### Source of Data

Mining of the scientific literature was conducted by using all 2,472,725 XML files from PubMed Central (commercial and non-commercial use), which consisted of information about each publication such as title, authors, DOI, abstract, full body content, and references.^41^ Clinical data for genetic mutations of HBV was collected from a total of 186 plasma samples from a Western European cohort of chronic HBV patients during the period from 1985 to 2012.^32^ HBV DNA was extracted from 200 *μ*L of plasma using the QIAamp MinElute Virus Kit (Qiagen) or the Roche MagNA Pure LC instrument according to the manufacturer’s instructions. These samples are stored at the Erasmus University Medical Center in Rotterdam, the Netherlands. The guidelines followed in the study were in accordance with the Declaration of Helsinki^42^ and the principles of Good Clinical Practice.^43^ The study was approved by the ethical review board of the Erasmus Medical Center, Rotterdam, the Netherlands. The dataset used to build the anti-HBV drugs’ vocabulary included 69 launched anti-HBV drugs obtained from the Cortellis Drug Discovery Intelligence database (CDDI), formerly known as Integrity.^44^ We extracted each drug’s code name, generic name, brand name, and drug name from the CDDI database to enrich this vocabulary, which gave 312 unique drug expressions.

### Mining the Scientific Literature

The workflow in Figure 1 was used to search through the sentences from journal articles in PubMed Central (PMC) that mentioned both an HBV drug and a mutation. In Step 2, we have used GNU parallel in order to select relevant literature that refer to ‘hepatitis b’ and/or ‘hbv’ (both keywords are case insensitive). Mutations used in Step 4 in Figure 1 were identified by using regular expressions for HBV mutations, as described in the next section. The output data from this workflow was later used to conduct the data analysis (*i.e*. Step 5 in Figure 1). Further details of Steps 1-5 are available at: https://github.com/angoto/HBV_Code.

**Figure 1.**
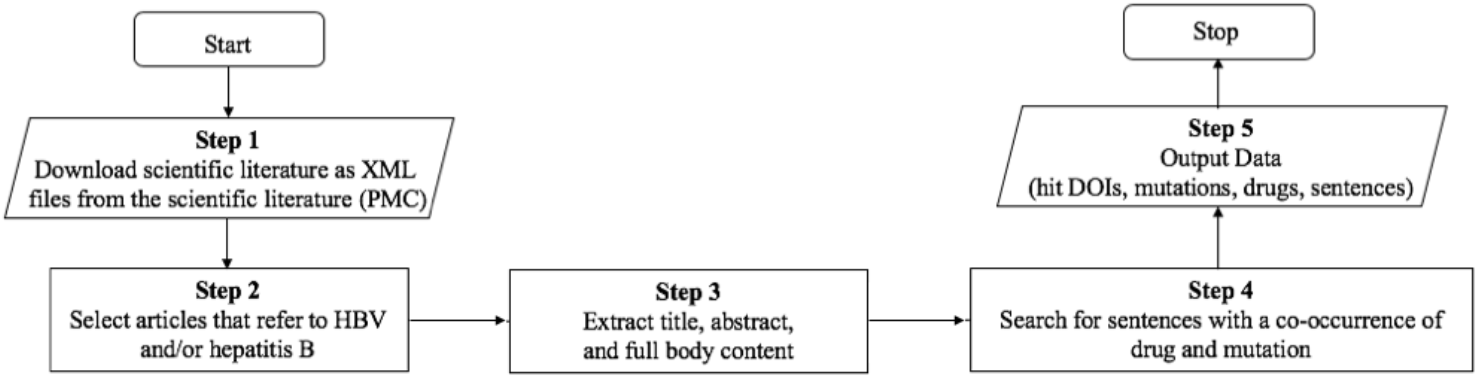
Workflow for mining the scientific literature.

### Regular Expressions for HBV Mutations

Mutations relevant to HBV were found from articles in PubMed Central by combining regular expressions.

### Use of Clinical Study Data

Six different pieces of data from the Western Europe cohorts described in Mueller-Breckenridge, *et al*.^32^ were used to conduct the analysis: sample, reference genome, gene, effect, amino acid variant, and nucleotide variant. “Sample” identifies the unique patients’ number used during the clinical study. Reference genome refers to a genotype-specific reference sequence (namely, GenBank accession numbers AF090842, AB033554, AB033556, AF121240, and AB032431, for genotypes A to E, respectively) that was used to map the outputs generated from quality-trimmed demultiplexed FASTQ reads.^32^ “Gene” indicates four known genes encoded by the genome: C, P, S, and X. “Effect” refers to whether a particular mutation was either an intragenic or a missense variant. For this report, we have only considered missense variants because intragenic variants from the literature were not associated with the clinical study’s phenotypes. “Amino acid variant” (*n* = 7,285) and “nucleotide variant” (*n* = 10,658) are two different ways to describe the same mutation, *i.e*. mutation in the amino acid (*e.g*. p.Pro156Ser) and nucleotide (*e.g*. c.314C>T) sequences, respectively. Although it is relatively straightforward to obtain HBV-related genetic mutations from the literature, it is difficult to map the position number of nucleotides to the position number of amino acids, and *vice-versa*. This is because we are unsure about the reference sequence that was used to report these mutations for each journal article in PubMed Central. In order to do this type of mapping with our current approach, we would need to manually go over each journal article that mentioned both an HBV drug and a mutation to conduct the translation of DNA sequences to amino acid sequences because an ambiguous inverse mapping would be impossible (with the exception of Tryptophan). Hypothetical clinical study data is available on the GitHub repository for this study^45^ to provide a better idea of the data structure used in the clinical study.

### Comparison of Scientific Literature and Clinical Study

The workflow in Figure 2 was used to compare mutations from the clinical study with the genetic mutations found in the scientific literature downloaded from PubMed Central after translating the mutations from the literature to the format used in the clinical study: amino acid (category *vi*) and nucleotide (category *ii*) variants (*n* = 4,214).

**Figure 2.**
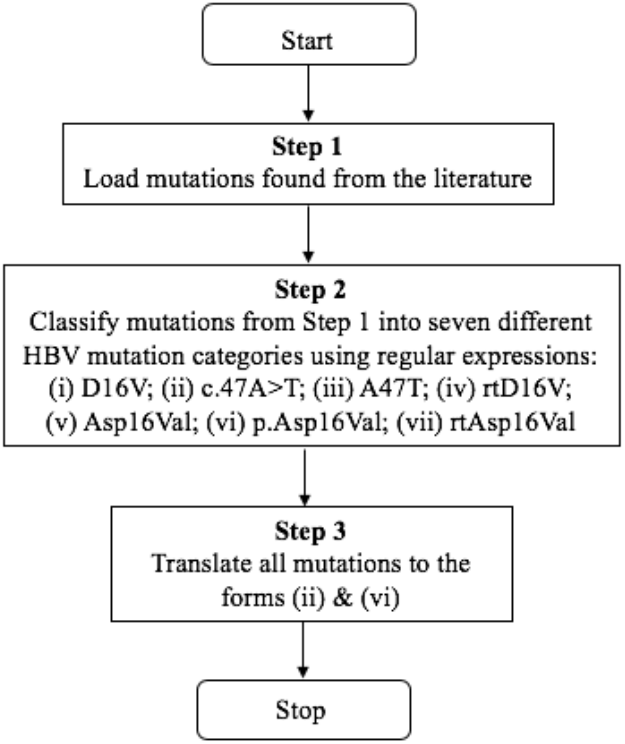
Normalization of DNA and amino acid mutations found from text mining the literature, to enable the direct comparison with clinical study mutations.

### Mutation Hotspot for Entecavir

We independently built our own models of entecavir bound with HBV RT and DNA, using the X-ray crystal structures of HIV RT also bound to entecavir and DNA, from PDB entries 5XN1 and 6IKA.^46,47^ Homology modelling was performed using SWISS-MODEL,^48^ and UniProt ID Q9WRJ9 residues 349-692 for the sequence of HBV RT genotype A.^49^ The resulting model underwent energy minimization using the open source build of PyMOL version 2.3.0,^50^ relaxing the entecavir ligand and all residues and bases within 4 Å of entecavir.

## Results

### Comparison between the Clinical Study and the Literature

The number of publications found to have a sentence co-occurrence of an approved HBV drug and a mutation in the same sentence was 30,686. There were 4,214 unique mutations mentioned in those sentences and, 7.5% of those, a total of 316 mutations (*i.e*. 254 amino acid and 62 nucleotide variants) were also found in the clinical data from Western Europe cohorts (*n* = 182 patients, 31,977 mutations). After analyzing the clinical data using solely those unique mutations that were common between the clinical study and the literature, we found that 180 out of 182 patients (corresponding to 1,750 amino acid and 302 nucleotide variants) had mutations that also appeared in the literature.

### Prevalence of Genetic Mutations in the Clinical Study

Figure 3 shows the ten most frequent amino acid (left) and nucleotide variants (right) in the clinical study that were also found in the literature. Their prevalence ranged between 1 and 52 patients for amino acid variants and between 1 and 78 patients for nucleotide variants. Figures S1-S2 show the frequency of mutations in the clinical data (*i.e*. number of patients reported to have each mutation type).

**Figure 3.**
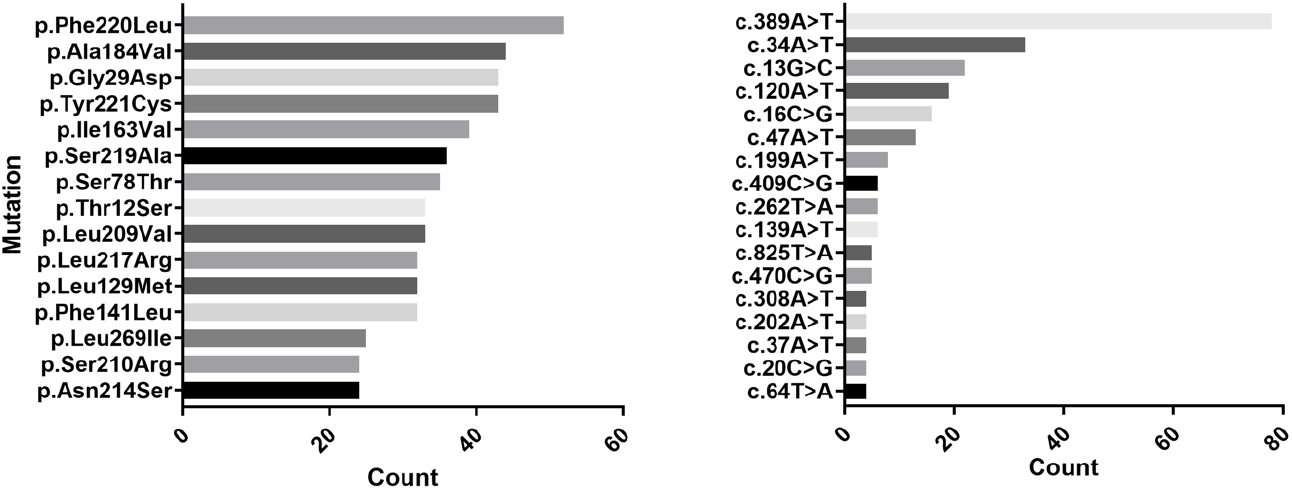
Top 10 most frequent amino acid mutations (left) and nucleotide mutations (right) in the clinical data that were also found in the literature.

### Mutation Likelihood of Patients in the Clinical Study

The 180 patients presented between 16 and 753 HBV mutations. Out of these mutations, between 1 and 35 appeared in the literature (Figure S3-1), with an average of 11.2 mutations and a median of 10 mutations. Figure S3-2 shows the patients in the clinical study who had the most mutations matching those found in the literature.

### Mutation Hotspots for HBV Genotypes in the Clinical Study

We identified amino acid mutations in the four genes X, P, C, and S, across five HBV genotypes, A, B, C, D, and E, that emerged during the clinical study. There were 182 patients in the clinical study, of which there were 56 patients with genotype A, 19 with genotype B, 43 with genotype C, 62 with genotype D, and 2 with genotype E. In order to create a mutation hotspot map for each of these genotypes (Figures S4-S8) where there was at least one report in the literature of that mutation, overlaps between the mutations from the literature and the clinical study mutations were identified. It should be noted that there are mutations in the patients that were present in the clinical study, but were not found in the analysis of the literature, and hence are not shown in these heatmaps.

We defined a “mutation hotspot” as a mutation with a count above the average for that particular gene and genotypes. For example, for HBV RT genotype A, mutation V214A had a count of 1 patient, while the average of the counts was 5.18 patients, so this was excluded from the map; while mutation L217R had a count of 30 patients, which was above the average for HBV RT genotype A, and thus was included as a mutation hotspot. For nucleotide variants, similar plots were made based on the genotypes (A-E), with mutations sorted in order of position number (Figures S9-S13). Figures 4–8 represent heatmaps of the mutational hotspots that we identified for eight different gene products (*i.e*. polymerase, reverse transcriptase, X, precore, core, PreS1, PreS2, and HBsAg) and nucleotide variants, with darker, more saturated colors indicating more mutations at that position in that genotype. The summary tables of hotspots for each HBV genotype for amino acid and nucleotide variants are shown in Figures S14-S18.

**Figure 4.**
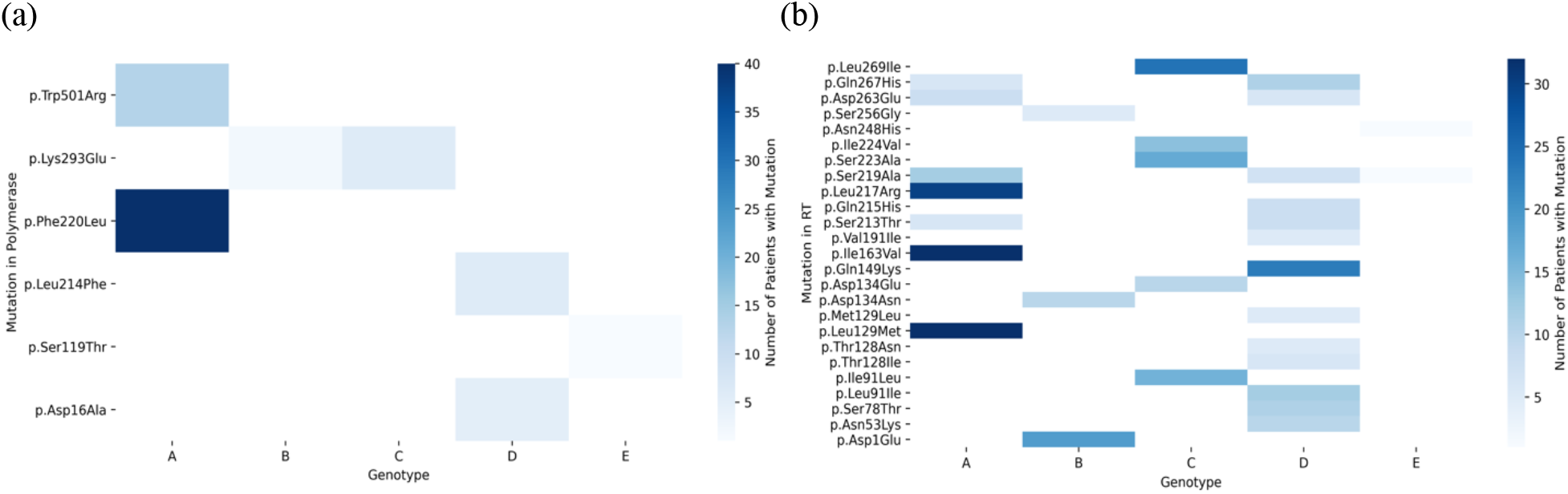
Heatmap of mutation hotspots for gene P in genotypes A-E: (a) HBV polymerase, and (b) HBV reverse transcriptase (RT). The white regions represent counts of mutations that are null, while dark blue indicates the largest number of mutations.

**Figure 5.**
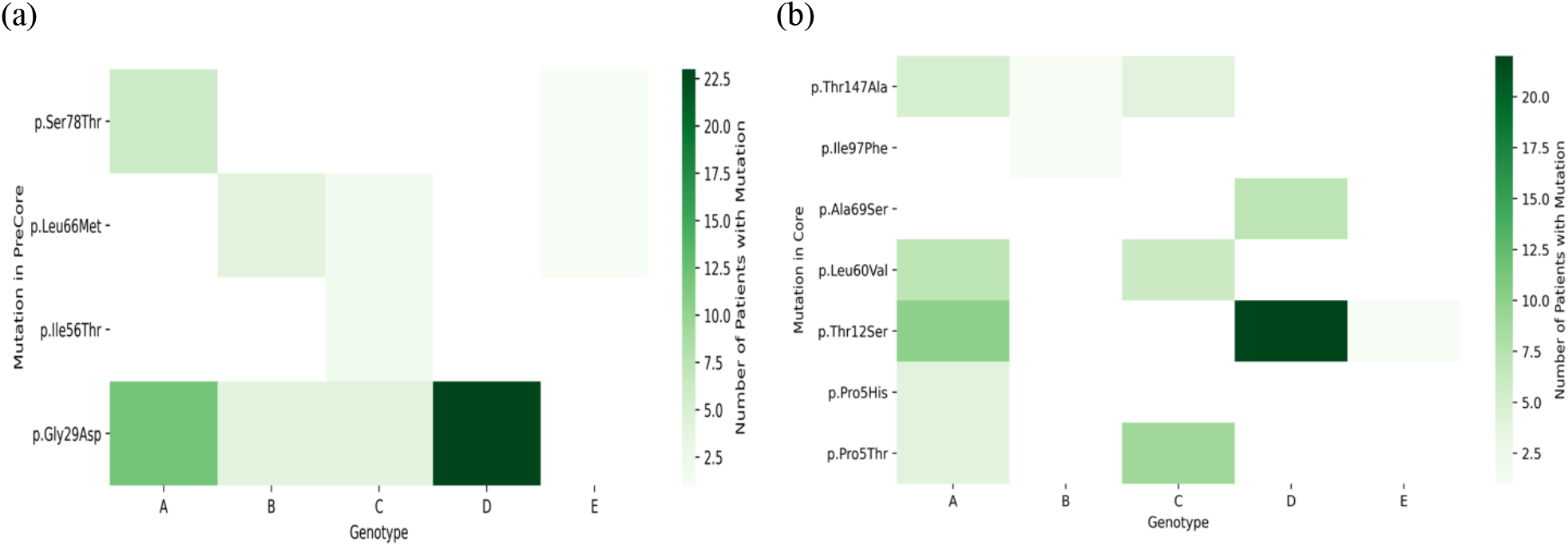
Heatmap of mutation hotspots for gene C in genotypes A-E: (a) HBV precore, and (b) HBV core. The white regions represent counts of mutations that are null, while dark green indicates the largest number of mutations.

**Figure 6.**
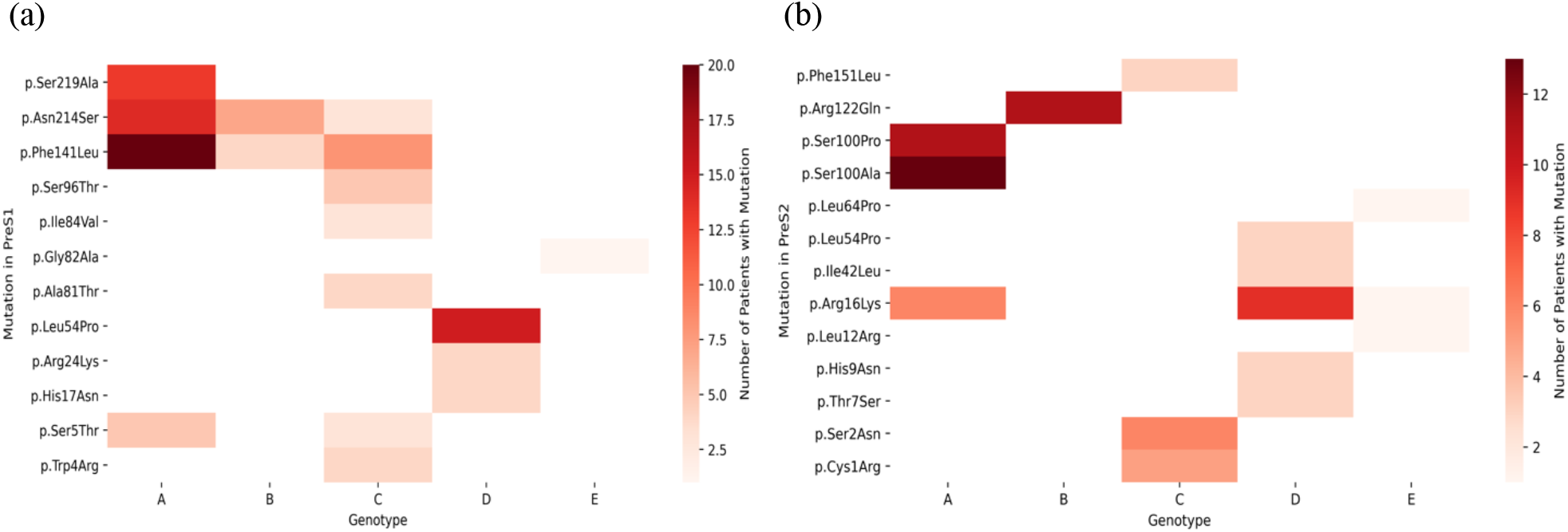
Heatmap of mutation hotspots for gene S in genotypes A-E: (a) HBV PreS1, and (b) HBV PreS2. The white regions represent counts of mutations that are null, while dark red indicates the largest number of mutations.

**Figure 7.**
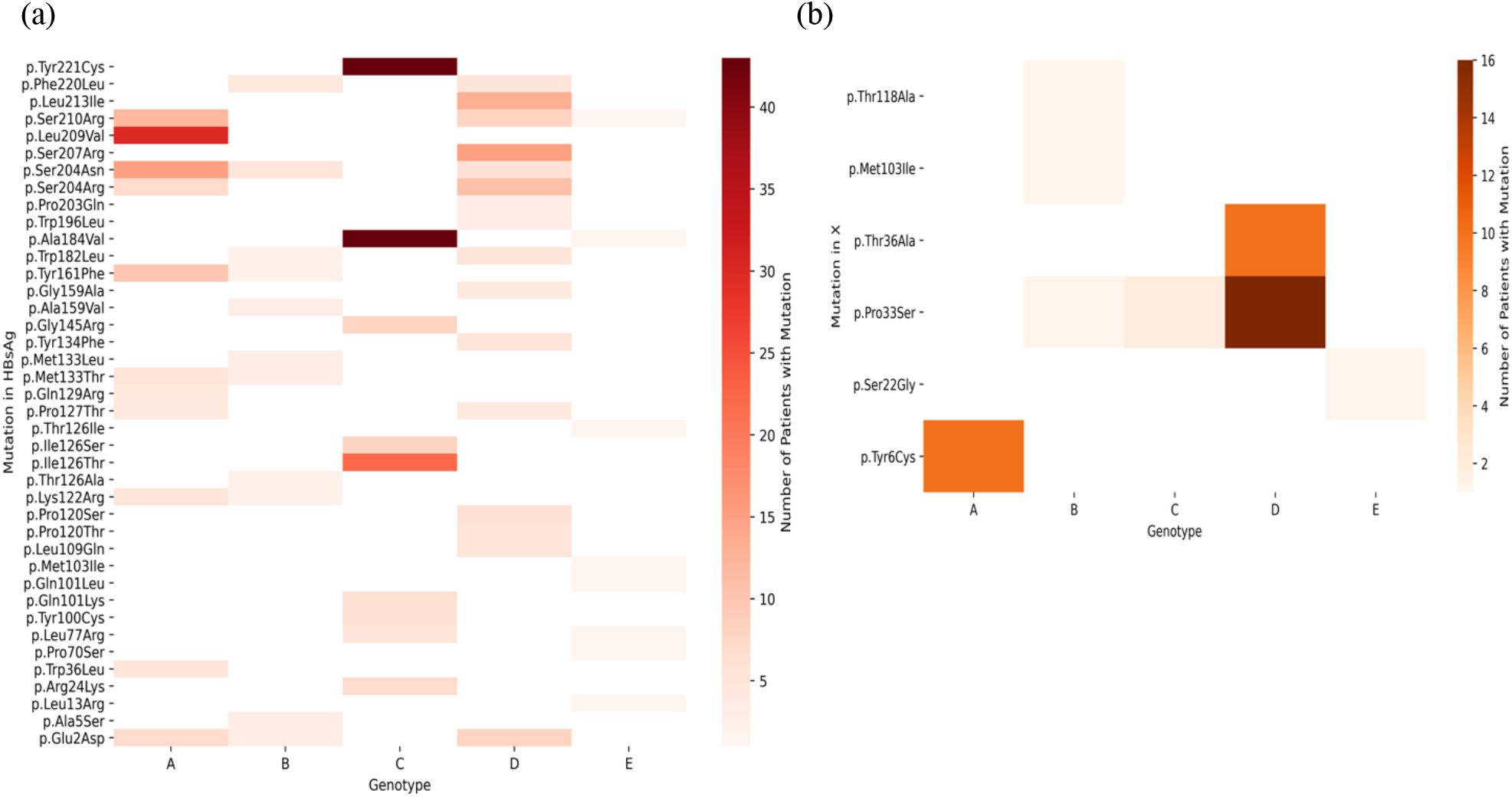
Heatmap of mutation hotspots for: (a) HBV HBsAg (gene S), and (b) HBV gene X in genotypes A-E. The white regions represent counts of mutations that are null, while dark red indicates the largest number of mutations.

**Figure 8.**
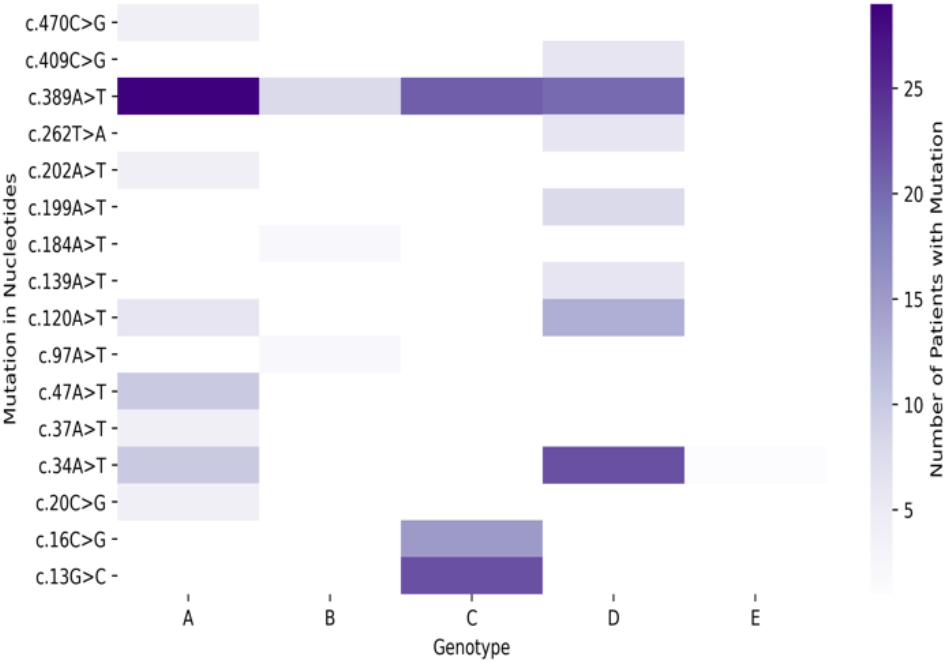
Heatmap of mutation hotspots for HBV nucleotide variants in genotypes A-E. The white regions represent counts of mutations that are null, while dark purple indicates the largest number of mutations.

Our analysis revealed there were “genotype-common” mutation hotspots, *i.e*. mutations with the same position number, in two or more genotypes (Figures 4–8), above the average count of patient. These were: polymerase at position K293; reverse transcriptase at positions S213, S219, D263, and Q267; PreS1 at positions S5, F141, and N214; PreS2 at position R16; HBsAg at positions E2, L77, K122, P127, M133, Y161, W182, A184, S204, S210, and F220; X at position P33; precore at positions G29, L66, and S78; core at positions P5, T12, L60, and T147; and nucleotide variants at positions 34A, 120A, and 389A.

Thus, although we were unable to discriminate HBV drug-resistant mutations unambiguously from other mutations, we were able to identify common mutations in the clinical study and the literature, across genotypes A-E. This can further our understanding of HBV mutations by highlighting mutations that may be potentially relevant to resistance.

### Mutations Occurring in Disulfide Bonds

We investigated the number of common mutations between the clinical study and the literature that arose from mutating wild type cysteine residues. Cysteine residues can be responsible for the formation of disulfide bonds, which play an important role in folding and stability of the proteins.^51^ The HBV envelope proteins, which corresponds to gene S, are known to form an intermolecular disulfide network through cysteine residues in the cysteine-rich antigenic loop that are located in positions 102 to 161.^52–54^ Hence, mutations for cysteine residues reported in Table 1 for positions C125 and C149 are in fact in regions of disulfide bonds. For gene C, the cysteine residue in position C48 is expected to form a disulfide bond with C149.^55^ We were unable to find any evidence of a disulfide bond involving residue C1 in gene S.

**Table 1.** Cysteine residues in genotypes A-E for common mutations between the clinical study and the literature with corresponding genes labelled in parentheses (*i.e*. gene S, and gene C).

| Genotype | Cysteine Residues |
| --- | --- |
| A | p.Cys149Arg (S), p.Cys48Gly (C) |
| B | p.Cys1Arg (S), p.Cys1Ser (S), p.Cys48Gly (C) |
| C | p.Cys149Arg (S), p.Cys1Arg (S) |
| D | p.Cys124Arg (S), p.Cys149Arg (S) |
| E | N/A |

Therefore, by comparing the literature and the clinical study, we were able to identify which gene, and in what particular position, mutations occurred at disulfide bond. We found mutations located at C124 and C149 in gene S to occur in regions of disulfide bonds, which are likely to have an effect towards destabilizing the HBV proteins. Table 1 shows the list of cysteine residues in each genotype, A-E, for mutations that were common between the clinical data and the literature.

### Mutation Hotspot for Entecavir

Based on a search conducted using DrugBank,^56^ one out of 69 approved drugs listed in the CDDI database had a modelling study published with a known target and binding site.^57^ By using our own models of entecavir bound with HBV RT and DNA, we identified which of these binding pocket residues were common between the clinical data and the scientific literature. For entecavir, we found common mutations located at I169, M204, and N238 for genotype A (within 12 Å of entecavir in our model).

Figure 9 shows the binding pocket of entecavir, together with the binding pocket residues within 5 Å of entecavir. The original modelling study by Langley, *et al*.^57^ used a sequence alignment between HIV and HBV reverse transcriptase to determine the most conserved domains, and they used the HIV reverse transcriptase DNA X-ray structure (PDB ID 1RTD)^58^ to build the HBV RT model.

**Figure 9.**
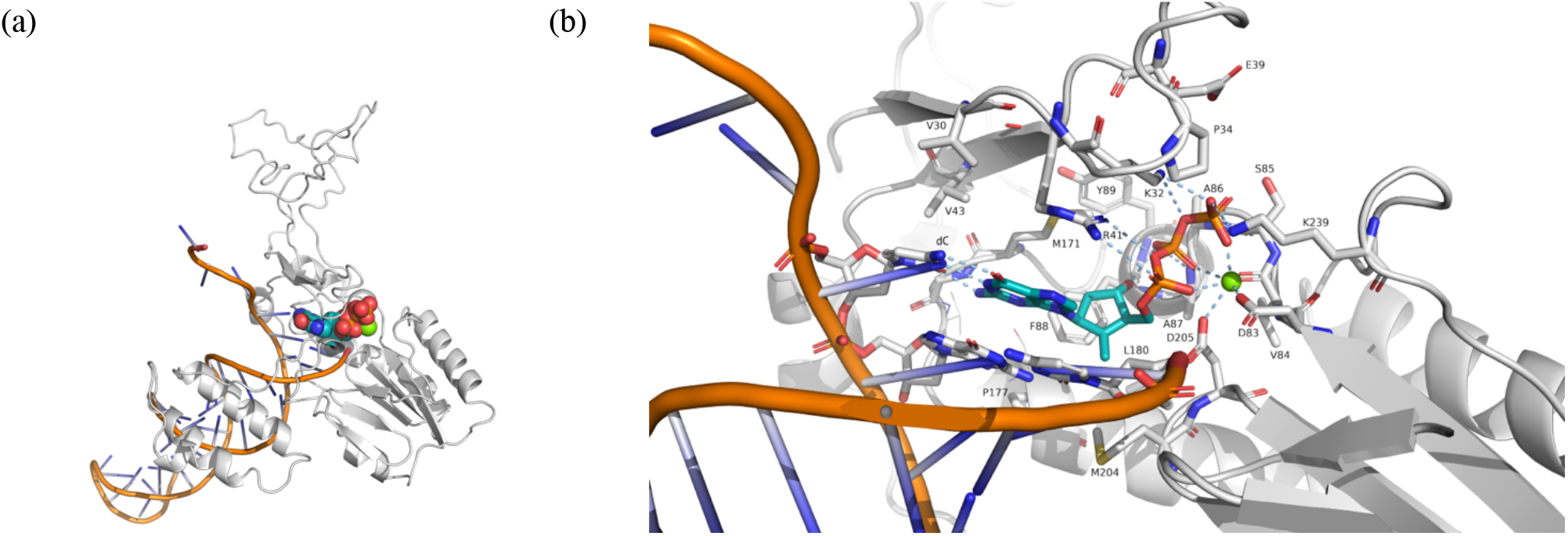
Model of HBV RT (genotype A) based on the HIV RT-DNA-entecavir complex X-ray crystal structure, PDB ID 5XN1.^46^ (a) Entecavir-triphosphate is shown as spheres with teal-colored carbon atoms, red oxygen, blue nitrogen, and orange phosphorus atoms. Entecavir is at the 3’ end of the DNA strand (orange cylinders). The secondary structure of the HBV RT model is shown as white coils (*α*-helices), white arrows (*β*-strands), and white loops. (b) Close-up view of the binding pocket, showing entecavir-triphosphate with teal-colored carbons, and a Mg^2+^ cation as a green sphere. Note that hydrogen bonds and metal bonds are shown as light blue dashed lines. The deoxyguanine (dG)-moiety of entecavir can be seen “base-pairing” with deoxycytosine (dC) in a second strand of DNA.

We also investigated the location of the mutational hotspots we identified for HBV RT in genotype A, and mapped these to our model. It can be seen from Figure 10 that the mutations occur throughout the structure.

**Figure 10.**
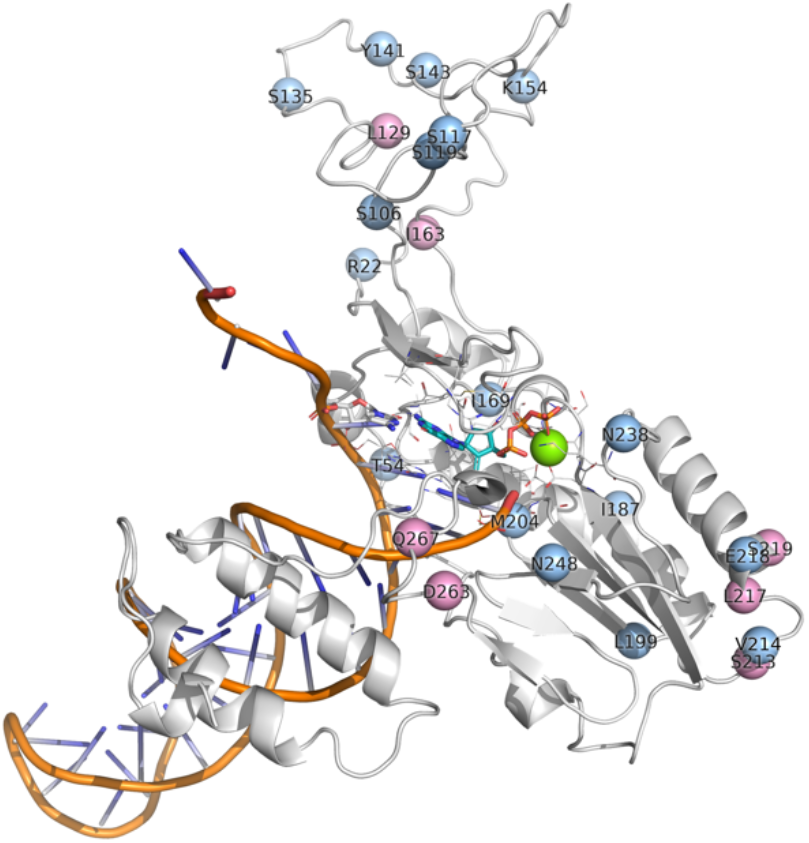
Locations of the mutations in HBV RT found both in patients in the clinical study, and in our analysis of the literature, shown as spheres placed at the *α* -carbon atom of the amino acid. The pink spheres are mutations with above-average counts of patients, and are at positions 129, 163, 213, 217, 219, 263, and 267; while the light blue spheres indicate positions of mutations below the average threshold. The secondary structure of our model of the HBV RT is shown as a white cartoon, while the DNA backbone is in orange. The location of the DNA, entecavir-triphosphate (shown with teal carbons), and Mg^2+^ (green sphere) was modelled on PDB ID 5XN1.^46^

## Discussion and Conclusions

This study showed the feasibility of integrating clinical and text mined viral mutation data and its potential to produce new insights on disease-relevant viral mutations. The focus of the study was on HBV but the methods are applicable to other viruses. While mutational hotspots can be identified by considering only the clinical study’s mutations, the difficulty associated with this process is that, even when new mutational hotspots are identified through the study, they are often disregarded because there are no literature reports available or there are not enough resources to comb the literature to confirm its role in resistance mutations. Thus, it is typical for clinical studies to report only those mutational hotpots that have been reported in the past. Hence, the methodology presented in this study serves as a bridge to fill in that gap by leveraging the full text of 2.47 million articles from PubMed Central to produce a landscape of disease-specific viral mutations. Although it is not possible for this method to confirm a direct correlation between the mutational hotspots and resistance mutations, it is able to provide new hypotheses that are potentially relevant to the resistance phenotype.

Our study identified known hotspots that were found in four genes, P, C, S, and X, in genotypes A to E, as shown in Figures 4–8. There are other HBV mutational hotspots that are known such as L180, which is a compensatory mutation for resistance to entecavir, lamivudine, and telbivudine;^59^ rt202 which causes resistance to entecavir in Asian population;^60^ and rt236 which is responsible for resistance to adefovir dipivoxil in Caucasian.^61^ Hence, the list of mutational hotspots that were identified in this study was not necessarily conclusive. In addition, two amino acid positions (*i.e*. C124 and C149) exhibited mutations in disulfide bonds, which are likely to impact the structure of HBV proteins. This finding is confirmatory since these regions of the disulfide bonds have already been identified in the past.^52–54^ Moreover, we identified a mutation in a position that was relevant to the binding site for the anti-HBV drug, entecavir. However, the I169 mutation, which is known to be the primary mutation responsible for entecavir’s resistance was found to be reported below the average threshold when comparing mutational hotspots between the clinical study and the literature.^62^ Therefore, further refinement of the definition of mutational hotpots used in this study may be necessary in the future.

One limitation of our approach lies in terms of the amount of data we collected from the literature. This resulted in an overall coverage of the number of genetic mutations reported in the literature against the common mutations of the literature and the clinical study of 7.5%. In order to further increase the coverage of genetic mutations data, literature sources could be increased to include repositories from publishers such as ScienceDirect^63^ and Springer Nature^64^. We could also obtain additional data from the PubMed Central literature by mining the information included in tables and figures. It is also important to be aware of errors that may arise during DNA sequencing and patient data collection during the clinical study.

In addition, we have not taken into account the rate of false positive and false negative identifications of mutations. An example of a false positive would be the following sentence: “*Some non-resistance-associated mutations of rtD134N (ranging from 20.33% to 74.63%), rtL145M (ranging from 2.83% to 78.82%*), *rtF151Y (ranging from 2.92% to 75.51%) and rtS223A (ranging from 5.77% to 18.44%) increased significantly with ADV monotherapy*, *then declined with the addition of LdT*”.^65^ In this particular sentence, it satisfies the criteria for co-occurrence of an HBV drug and a mutation, but it is in fact not referring to resistance mutations. False positives could also be considered hypothetical mutations, such as the one described in the sentence: “*The lamivudine resistance-linked G529A (rtD134N) site in HBV was found to be associated with HCC outcomes, which implied potential correlation between resistance to the anti-HBV nucleoside analog lamivudine and HCC prognosis*”.^66^ Furthermore, it is possible that some drug-resistant mutants involve multiple locations, so in the future we must consider such combinatorial possibilities as well.

To improve the methods used here, additional text mining strategies could be used to improve the extraction, such as machine learning algorithms, thus increasing our confidence that those sentences describe HBV-related information. For example, the following sentence refers to two mutations, M204I and M204V, which are not captured by simple regular expressions: “*LVDr is well characterized and arises through replacement of M204 within the YMDD motif of the HBV RT with isoleucine or valine, with or without the adaptive change L180M*”.^49^ By performing the literature mining in this manner, we could be able to develop a more powerful text mining tool that would allow us to identify a more comprehensive set of resistance mutations. However, it is important to note that, while more-recently published mutation-extraction approaches are machine-learning based,^67,68^ these require costly gold standard corpora, which hinders their re-application to different viral species, each with its own notation particularities. Thus, the development of machine-learning algorithms may lead to improved performance but could result in more restricted application to a particular disease.

## Availability of Data and Materials

The code and supplementary information (Figures S1-S18) is available at https://github.com/angoto/HBV_Code.

## Acknowledgments

We thank the Roche Innovation Center Basel for their computational support. A.G and G.M.M thank the EPSRC and MRC Centre for Doctoral Training (CDT) in Systems Approaches to Biomedical Science (EP/L016044/1). A.G thanks the Clarendon and Oxford-Kobe Scholarships. G.M.M thanks the EPSRC CDT in Sustainable Approaches to Biomedical Science: Responsible and Reproducible Research – SABS:R^3^ (EP/S024093/1).

## References

1. Polaris Observatory Collaborators. Global prevalence, treatment, and prevention of hepatitis B virus infection in 2016: a modelling study. Lancet Gastroenterol Hepatol. 2018;3(6):383–403.

2. Sarin SK, Kumar M, Lau GK, et al. Asian-pacific clinical practice guidelines on the management of hepatitis B: a 2015 update. Hepatol Int. 2016;10(1):1–98.

3. World Health Organization. Hepatitis B [Internet]. World Health Organization; 2019 [Accessed 2019 Sep 12]. Available from: https://www.who.int/en/news-room/fact-sheets/detail/hepatitis-b.

4. Lok ASF, McMahon BJ, Brown Jr RS, et al. Antiviral therapy for chronic hepatitis B viral infection in adults: A systematic review and meta-analysis. Hepatology. 2016;63(1):284–306.

5. Spradling P, Hu D, McMahon BJ. Epidemiology and prevention. In Viral hepatitis, 4th ed. (ed. Thomas H, et al.), Wiley, Hoboken, NJ. 2013. pp. 81–95.

6. Tang LSY, Covert E, Wilson E, Kottilil S. Chronic Hepatitis B Infect A Review. JAMA 2018;319(17):1802–1813.

7. Liang TJ. Hepatitis B: The virus and disease. Hepatology. 2009;49(5);S13–S21.

8. Kay A, Zoulim F. Hepatitis B virus genetic variability and evolution. Virus Res. 2007;127(2):164–167.

9. Buti M, Rodriguez-Frias F, Jardi R, Esteban R. Hepatitis B virus genome variability and disease progression: the impact of pre-core mutations and HBV genotypes. J. Clin Virol. 2005;34:S79–S82.

10. Beck J, Nassal M. Hepatitis B virus replication. World J Gastroenterol. 2007;13(1):48–64.

11. Orito E, Mizokami M, Ina Y, et al. Host-independent evolution and a genetic classification of the hepadnavirus family based on nucleotide sequences. Proc Natl Acad Sci USA. 1989;86(18):7059–7062.

12. Cross JC, Wen P, Rutter WJ. Transactivation by hepatitis B virus X protein is promiscuous and dependent on mitogen-activated cellular serine/threonine kinases. Proc Natl Acad Sci USA. 1993;90(17):8078–8082.

13. Hu Z, Zhang Z, Kim JW, Huang Y, Liang TJ. Altered proteolysis and global gene expression in hepatitis B virus X transgenic mouse liver. J Virol. 2006;80(3):1405–1413.

14. Milich D, Liang TJ. Exploring the biological basis of hepatitis B e antigen in hepatitis B virus infection. Hepatology. 2003;38(5):1075–1086.

15. Zhang Z, Torii N, Hu Z, Jacob J, Liang TJ. X-deficient woodchuck hepatitis virus mutants behave like attenuated viruses and induce protective immunity in vivo. J Clin Invest. 2001;108(10):1523–1531.

16. Lamivudine. DrugBank [Internet]. DrugBank [Accessed 2019 Sep 23]. Available from: https://www.drugbank.ca/drugs/DB00709.

17. Adefovir dipivoxil. DrugBank [Internet]. DrugBank [Accessed 2019 Sep 23]. Available from: https://www.drugbank.ca/drugs/DB00718.

18. Dienstag JL, Schiff ER, Wright TL, et al. Lamivudine as initial treatment for chronic hepatitis B in the United States. N Engl J Med. 1999;341(17):1256–1263.

19. Chan HL, Wang H, Niu J, et al. Two-year lamivudine treatment for hepatitis B e antigen-negative chronic hepatitis B. a double-blind, placebo-controlled trial. Antivir Ther. 2007;12(3):345–353.

20. Marcellin P, Chang T, Lim SGL, et al. Long-term efficacy and safety of adefovir dipivoxil for the treatment of hepatitis B e antigen-positive chronic hepatitis B. Hepatology. 2008;48(3):750–758.

21. Hadziyannis SJ, Tassopoulos NC, Heathcote EJ, et al. Long-term therapy with adefovir dipivoxil for HBeAg-negative chronic hepatitis B for up to 5 years. Gastroenterology. 2006;131(6):1743–1751.

22. Entecavir. DrugBank [Internet]. DrugBank [Accessed 2019 Sep 23]. Available from: https://www.drugbank.ca/drugs/DB00442.

23. Tenofovir disoproxil [Internet]. DrugBank [Accessed 2019 Sep 23]. Available from: https://www.drugbank.ca/drugs/DB00300.

24. Tenney DJ, Rose RE, Baldick CJ, et al. Long-term monitoring shows hepatitis B virus resistance to entecavir in nucleoside-naïve patients is rare through 5 years of therapy. Hepatology. 2009;49(5):1503–1514.

25. Buti M, Tsai N, Petersen J, et al. Seven-Year Efficacy and Safety of Treatment with Tenofovir Disoproxil Fumarate for Chronic Hepatitis B Virus Infection. Dig Dis Sci. 2015;60(5):1457–1464.

26. Bui QC, Nualláin BO, Boucher CA, & Sloot PM. Extracting causal relations on HIV drug resistance from literature. BMC Bioinformatics 2010;11:101.

27. Khalid Z, and Sezerman OU. ZK (2017)/ DrugResist 2.0: A TextMiner to extract semantic relations of drug resistance from PubMed. J Biomed Inform. 2017;69:93–98.

28. Naderi N, & Witte R. Automated extraction and semantic analysis of mutation impacts from the biomedical literature. BMC Genomics 2012;13:Suppl 4: S10.

29. Roberts K, Alam T, Bedrick S, Demner-Fushman D, Lo K, Soboroff I, Voorhees E, Wang LL, Hersh WR. TREC-COVID: rationale and structure of an information retrieval shared task for COVID-19. J Am Med Inform Assoc. 2020;27(9):1431–1436.

30. Davey NE, Satagopam VP, Santiago-Mozos S, Villacorta-Martin C, Bharat TA, Schneider R, Briggs JA. The HIV mutation browser: a resource for human immunodeficiency virus mutagenesis and polymorphism data. PLoS Comput Biol. 2014;10(12):e1003951.

31. Wang Y, Tong Y, Zhang Z, Zheng R, Huang D, Yang J, Zong H, Tan F, Zhang X. ViMIC: A Database of Human Disease-related Virus Mutations, Integration Sites and Cis-effects. bioRxiv. 29 Oct 2020.

32. Mueller-Breckenridge AJ, Garcia-Alcalde F, Wildum S, et al. Machine-learning based patient classification using Hepatitis B virus full-length genome quasispecies from Asian and European cohorts. Sci Rep. 2019;9(1):18892.

33. Caporaso JG, Baumgartner WA Jr, Randolph DA, Cohen KB, Hunter L. MutationFinder: a high-performance system for extracting point mutation mentions from text. Bioinformatics. 2007;23(14):1862–1865.

34. Doughty E, Kertesz-Farkas A, Bodenreider O, Thompson G, Adadey A, Peterson T, Kann MG. Toward an automatic method for extracting cancer- and other disease-related point mutations from the biomedical literature. Bioinformatics. 2011;27(3):408–415.

35. Thomas P, Rocktäschel T, Hakenberg J, Lichtblau Y, Leser U. SETH detects and normalizes genetic variants in text. Bioinformatics. 2016;32(18):2883–2885.

36. Hepatitis B Virus [Internet]. NCBI [Accessed 2021 June 24]. Available from: https://www.ncbi.nlm.nih.gov/genome/?term=hbv.

37. HBVdb [Internet]. [Accessed 2021 June 24]. Available from: https://hbvdb.lyon.inserm.fr/HBVdb/HBVdbAbout.

38. SeqHepB [Internet]. [Accessed 2021 June 24]. Available from: https://www.seqhepb.com/.

39. euresis network [Internet]. [Accessed 2021 June 24]. Available from: https://www.euresist.org/.

40. Python. xml.etree.ElementTree - The Element Tree XML API [Internet]. Python [Accessed 2019 Sep 13]. Available from: https://docs.python.org/3/library/xml.etree.elementtree.html.

41. NCBI. PMC [Internet]. National Center for Biotechnology Information: U.S. National Library of Medicine [Downloaded 2019 July 26, Accessed 2019 Sep 11]. Available from: https://www.ncbi.nlm.nih.gov/pmc/.

42. World Medical Association. World Medical Association Declaration of Helsinki. JAMA. 2013;310(20):2191–2194.

43. Vijayananthan A, Nawawi O. The importance of Good Clinical Practice guidelines and its role in clinical trials. Biomed Imaging Interv J. 2008;4(1):e5.

44. Clarivate Analytics. Integrity [Internet]. Clarivate Analytics [Accessed 2019 Sep 11]. Available from: https://integrity.clarivate.com//integrity/xmlxsl/.

45. GitHub. HBV_Code [Internet]. GitHub [Accessed 2019 Sep 20]. Available from: https://github.com/angoto/HBV_Code.

46. PDB ID: 5XN1. Yasutake Y, Hattori, SI, Hayashi, H, et al. HIV-1 with HBV-associated Q151M substitution in RT becomes highly susceptible to entecavir: structural insights into HBV-RT inhibition by entecavir. Sci Rep. 2018;8:1624–1624. DOI:10.1038/s41598-018-19602-9.

47. PDB ID: 6IKA. Yasutake Y, Hattori SI, Tamura N, et al. Active-site deformation in the structure of HIV-1 RT with HBV-associated septuple amino acid substitutions rationalizes the differential susceptibility of HIV-1 and HBV against 4’-modified nucleoside RT inhibitors. Biochem Biophys Res Commun 2019;509:943–948. DOI: 10.1016/j.bbrc.2019.01.026.

48. Bienert S, Waterhouse A, de Beer TAP, et al. The SWISS-MODEL Repository – new features and functionality. Nucleic Acids Res. 2017;45(D1):D313–D319.

49. UniProtKB - Q9WRJ9 (Q9WRJ9_HBV) [Internet]. UniProt [Accessed 2020 Feb 5]. Available from: https://www.uniprot.org/uniprot/Q9WRJ9.

50. The PyMOL Molecular Graphics System, Version 2.3.0 Schrödinger, LLC.

51. Sevier CS, Kaiser CA. Formation and transfer of disulfide bonds in living cells. Nat Rev Mol Cell Biol. 2002; 3(11):836–847.

52. Nassal M, Rieger A, Steinau O. Topological analysis of the hepatitis B virus core particle by cysteine-cysteine cross-linking. J Mol Biol. 1992; 225(4):1013–1025.

53. Mangold CM, Streeck RE. Mutational analysis of the cysteine residues in the hepatitis B virus small envelope protein. J Virol. 1993;67(8):4588–4597.

54. Mangold CM, Unckell F, Werr M, Streeck RE. Analysis of intermolecular disulfide bonds and free sulfhydryl groups in hepatitis B surface antigen particles. Arch Virol. 1997;142(11):2257–2267.

55. Mangold CM, Unckell F, Werr M, Streeck RE. Secretion and antigenicity of hepatitis B virus small envelope proteins lacking cysteines in the major antigenic region. Virology. 1995;211(2):535–543.

56. DrugBank [Internet]. DrugBank [Accessed 2019 Sep 12]. Available from: https://www.drugbank.ca.

57. Langley DR, Walsh AW, Baldick CJ, et al. Inhibition of hepatitis B virus polymerase by entecavir. J. Virol. 2007;81(8):3992–4001.

58. PDB ID: 1RTD. Huang H, Chopra R, Verdine GL, Harrison SC. Structure of a covalently trapped catalytic complex of HIV-1 reverse transcriptase: implications for drug resistance. Science 1998;282:1669–1675. DOI: 10.1126/science.282.5394.1669.

59. C. L. Lai, N. Leung, W. K. Teo et al. A 1-year trial of telbivudine, lamivudine, and the combination in patient with hepatitis B e antigen-positive chronic hepatitis B. Gastroenterology. 2005;129(2):528–536.

60. H. S. Kim, H. J. Yim, M. K. Jang et al. Management of entecavir-resistant chronic hepatitis B with adefovir-based combination therapies. World J. Gastroenterol. 2015;21(38):10874–10882.

61. P. Angus, R. Vaughan, S. Xiong et al. Resistance to adefovir dipivoxil therapy associated with the selection of a novel mutation in the HBV polymerase. Gastroenterology. 2003;125(2):292–297.

62. A. S. Lok and B. J. McMahon. Chronic hepatitis B: update of recommendations. Hepatology. 2004;39(3):857–861.

63. ScienceDirect [Internet]. ELSEVIER, ScienceDirect [Accessed 2019 Sep 19]. Available from: https://www.sciencedirect.com/.

64. Springer Nature [Internet]. Springer Nature [Accessed 2019 Sep 19]. Available from: https://www.springernature.com/gp.

65. Zhang X, Li M, Xi H, et al. Pre-existing mutations related to tenofovir in chronic hepatitis B patients with longterm nucleos(t)ide analogue drugs treatment by ultra-deep pyrosequencing. Oncotarget. 2016;7(43):70264–70275.

66. Lin C, Chien R, Hu C, Lai M, Yeh C. Identification of hepatitis B virus rtS117F substitution as a compensatory mutation for rtM204I during lamivudine therapy. J Antimicrob Chemother. 2012;67(1):39–48.

67. Cejuela JM, Bojchevski A, Uhlig C, Bekmukhametov R, Kumar Karn S, Mahmuti S, Baghudana A, Dubey A, Satagopam VP, Rost B. nala: text mining natural language mutation mentions. Bioinformatics. 2017;33(12):1852–1858.

68. Wei CH, Phan L, Feltz J, Maiti R, Hefferon T, Lu Z. tmVar 2.0: integrating genomic variant information from literature with dbSNP and ClinVar for precision medicine. Bioinformatics. 2018;34(1):80–87.

